# VREX: An Open-Source Toolbox for Creating 3D Virtual Reality Experiments

**DOI:** 10.1101/105858

**Authors:** Madis Vasser, Markus Kängsepp, Murad Magomedkerimov, Kälver Kilvits, Vladislav Stafinjak, Taavi Kivisik, Raul Vicente, Jaan Aru

**Author notes:** corresponding author: Madis Vasser.

## Abstract

**Background:** We present VREX, a free open-source Unity toolbox for virtual reality research in the fields of experimental psychology and neuroscience.

**Results:** Different study protocols about perception, attention, cognition and memory can be constructed using the toolbox. VREX provides a procedural generation of (interconnected) rooms that can be automatically furnished with a click of a button. VREX includes a menu system for creating and storing experiments with different stages. Researchers can combine different rooms and environments to perform end-to-end experiments including different testing situations and data collection. For fine-tuned control VREX also comes with an editor where all the objects in the virtual room can be manually placed and adjusted in the 3D world.

**Conclusions:** VREX simplifies the generation and setup of complicated VR scenes and experiments for researchers. VREX can be downloaded and easily installed from vrex.mozello.com

## Introduction

Conducting both ecologically valid and at the same time highly controlled psychological and neuroscientific experiments is a notoriously difficult task, preventing it from widespread use [1]. Controlling or manipulating every variable in real life is often not possible (e.g. making a couch silently disappear in an instance), too expensive (constructing a special building just for an experiment) or even dangerous (confronting someone with a hungry lion). Hence, researchers commonly settle to present 2D images on a computer screen. Obviously such conditions lack many real-life features and make it questionable how much of the cognition happening in the natural world can be captured in these simplified model environments [2].

However, 3D virtual reality environments are starting to provide a reasonable compromise between the expensive real-world and simplified computer screen experiments [2,3,4]. Head mounted displays (HMD) can render computer-generated quasi-realistic natural scenes, giving the experimenter more control over details of the environment, and allow near-perfect reproduction of the experimental setting between participants [5]. As the virtual environment inside a HMD is projected spherically all around the person when turning one’s head, this approach also gives more freedom of movement to the study participant, who is no longer confined to look only in a single narrow direction towards the computer monitor. Various studies can profit from a virtual reality implementation, as opposed to the "standard" computer screen version. Such topics include spatial navigation, perception and motor control [4]. Modern VR systems are capable of high level of immersion and the feeling of presence, as wider field of view, precise head tracking, improved visuals and audio have all been shown to increase participants’ subjective illusion of actually being in the virtual world [6]. Here, presence refers to the perception of one's surrounding as mediated by both automatic and controlled mental processes, an experience of a different reality [7]. High level of presence in a virtual study environment might yield stronger cognitive ethology [8], resembling the high perceptual and computational demands present in real life behaviors [4,9].

Recently launched consumer grade VR headsets such as the Oculus Rift and HTC Vive allow 360 degree optical tracking and up to 16 square meters of movement space, while at the same time being affordable. These hardware breakthroughs in the last few years have made VR technology available for almost any lab studying experimental psychology. Also, major game engines such as Unity (Unity Technologies) and Unreal Engine (Epic Games, Inc) now have built-in VR support straight out of the box. However, this new research paradigm requires specialized knowledge in software and hardware technology in order to create immersive and presence-inducing virtual realities. In particular, knowledge of 3D modeling and texturing, game engine logic and scripting are all needed.

Many psychology labs still lack these competences. The primary aim of the Virtual Reality Experiments (VREX) Toolbox is to help psychology researchers easily create experiments for virtual reality setups by providing an open-source software suites as an Unity add-on and standalone version, documentation and a web platform. Next we present related works, detailed descriptions of toolbox features and two possible use cases.

## Why VREX: Related work

There are a wide variety of Unity add-ons assisting the generation of interactive virtual worlds, such as Playmaker [10], Adventure Creator [11] and ProBuilder [12] to name a few. Yet these toolboxes are very general-purpose. There also exists some software applications similar to VREX in terms of simplifying the creation of VR experiments for psychological research, e.g. MazeSuite [13] and WorldViz Vizard [14]. The list of compared software is not comprehensive and here we briefly describe only two of them with key differences to VREX.

MazeSuite is a free toolbox that allows easy creation of connected 3D corridors. It enables researchers to perform spatial and navigational behavior experiments within interactive and extendable 3D virtual environments [13]. Although the user can design mazes by hand and fill them with objects, it is difficult to achieve the look and feel of a regular apartment. This is where VREX differs, having been designed for indoor experiments in mind from the beginning. Another noticeable difference is that MazeSuite runs as a standalone program, while VREX can be embedded inside Unity Game Engine, allowing for more powerful features, higher visual quality and faster code iterations in our experience.

WorldViz Vizard gives researchers the tools to create and conduct complex VR-based experiments. Researchers of any background can rapidly develop their own virtual environments and author complex interactions between environment, devices, and participants [14]. Although Vizard is visually advanced, this comes at a price of the licence fee to remove time restrictions and prominent watermarks. VREX matches the graphical quality of Vizard with the power of Unity 5 game engine, while staying open source and free of charge (Unity license fees may apply for publishing).

As any software matures, more features tend to be added by the developers. This in term means more complex interfaces that might confuse the novice user. The advantage of VREX is the narrow focus to specific types of experiments, allowing for clear design and simple workflow.

## Implementation

## Overall logic

The basic elements of the toolbox are environments consisting of single or multiple rooms. Environments can be sequenced to create experiments. Special pre-made environments can be added for displaying instructions to the participant or administering tests. Environments can be grouped together as ordered or randomized trial blocks. Thus an experiment can start with a text environment as instructions for the participant, then present procedurally generated rooms in a random order and end with a test block to gather responses. A typical pipeline for an experiment can be seen on figure 1.

**Figure 1.**
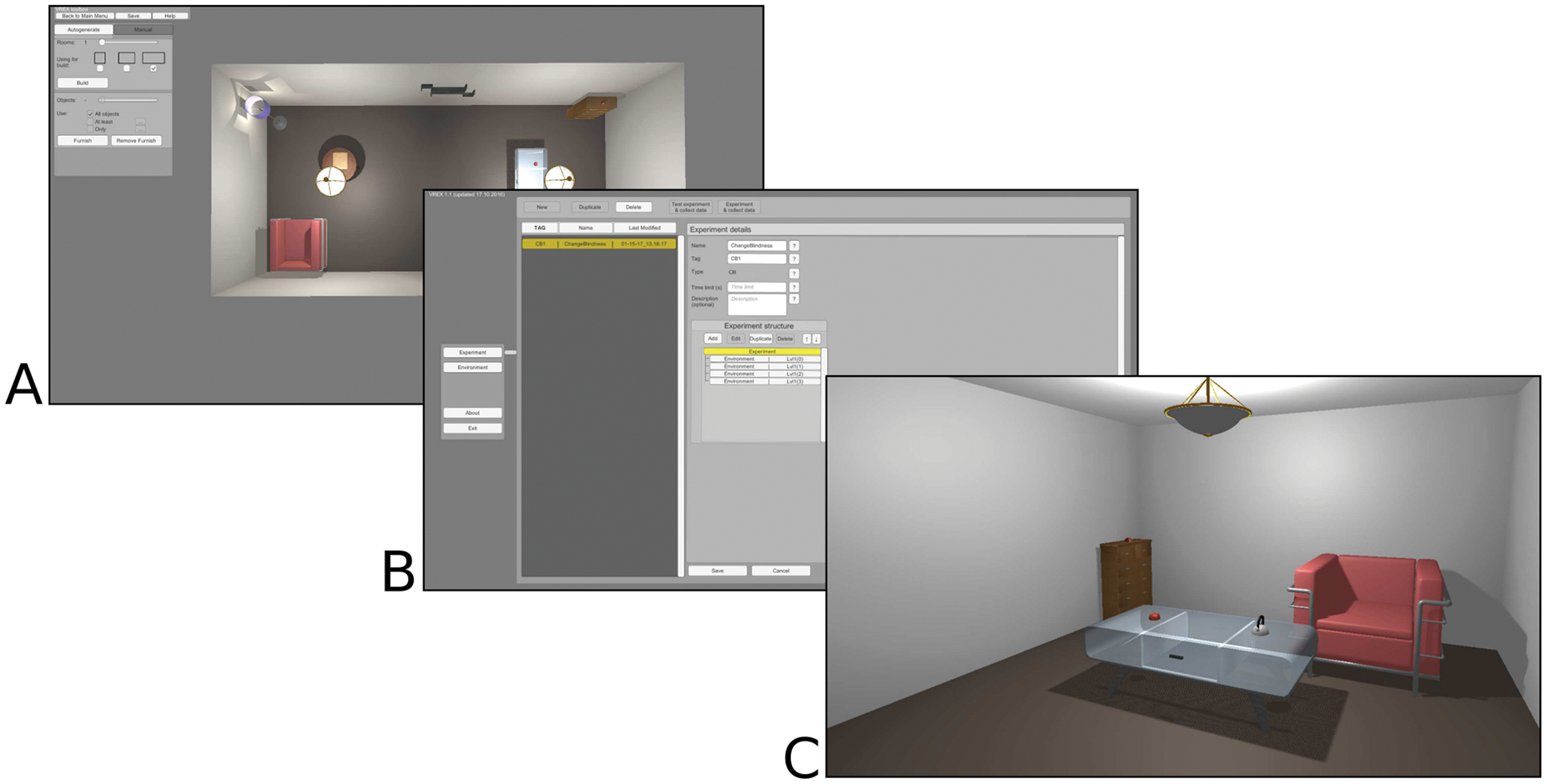
A typical pipeline for an experiment. Creating the environments (A), building the experiment structure (B) and running the study in VR (C)

## Graphical User Interface

Reading through and modifying C# code can be a taunting task. While Unity natively displays public variables in the inspector window, navigating a long array of options quickly gets overwhelming. Also, when building a standalone version, access to variables via user interface is lost. For these reasons VREX provides a separate graphical user interface inside the toolbox to give the user intuitive access to common operations within the program (figure 2). The simple menus allow creating and modifying environments and build experimental plans with different stages.

**Figure 2.**
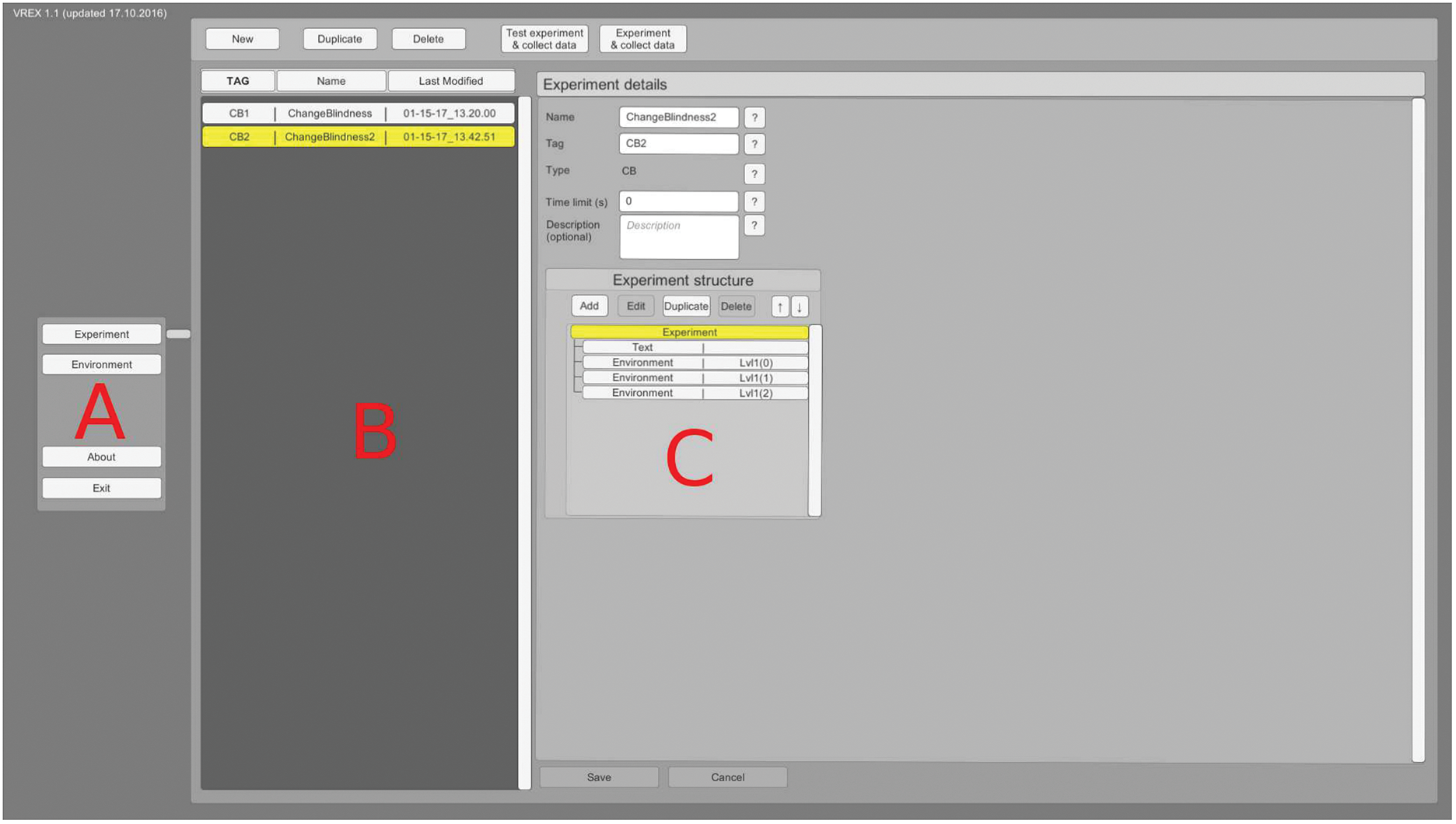
VREX menu layout. Main menu (A), list of created experiments (B) and the structure of a particular experiment (C).

## Creating environments

An environment is used as the main building block for experiments. Each environment consists of one or more rooms and may be populated with objects. The user can either create an environment from scratch or duplicate or modify an already existing environment. Starting from a blank scene, it is possible to either autogenerate the environment or combine rooms one-by-one manually (figure 3). The default option is autogeneration, as this feature saves time and provides an unique environment layout every time. For automatic generation of connected rooms it is mandatory to specify the number of desired rooms (up to 10). The user can also choose the dimensions of the rooms. Due to the algorithms employed, the rooms are either square (1x1), or rectangular (1x1.5 or 1x2). Autogeneration combines the rooms in a way that doorways connect and the geometry avoids overlapping. Currently VREX is confined to generating indoors environments.

**Figure 3.**
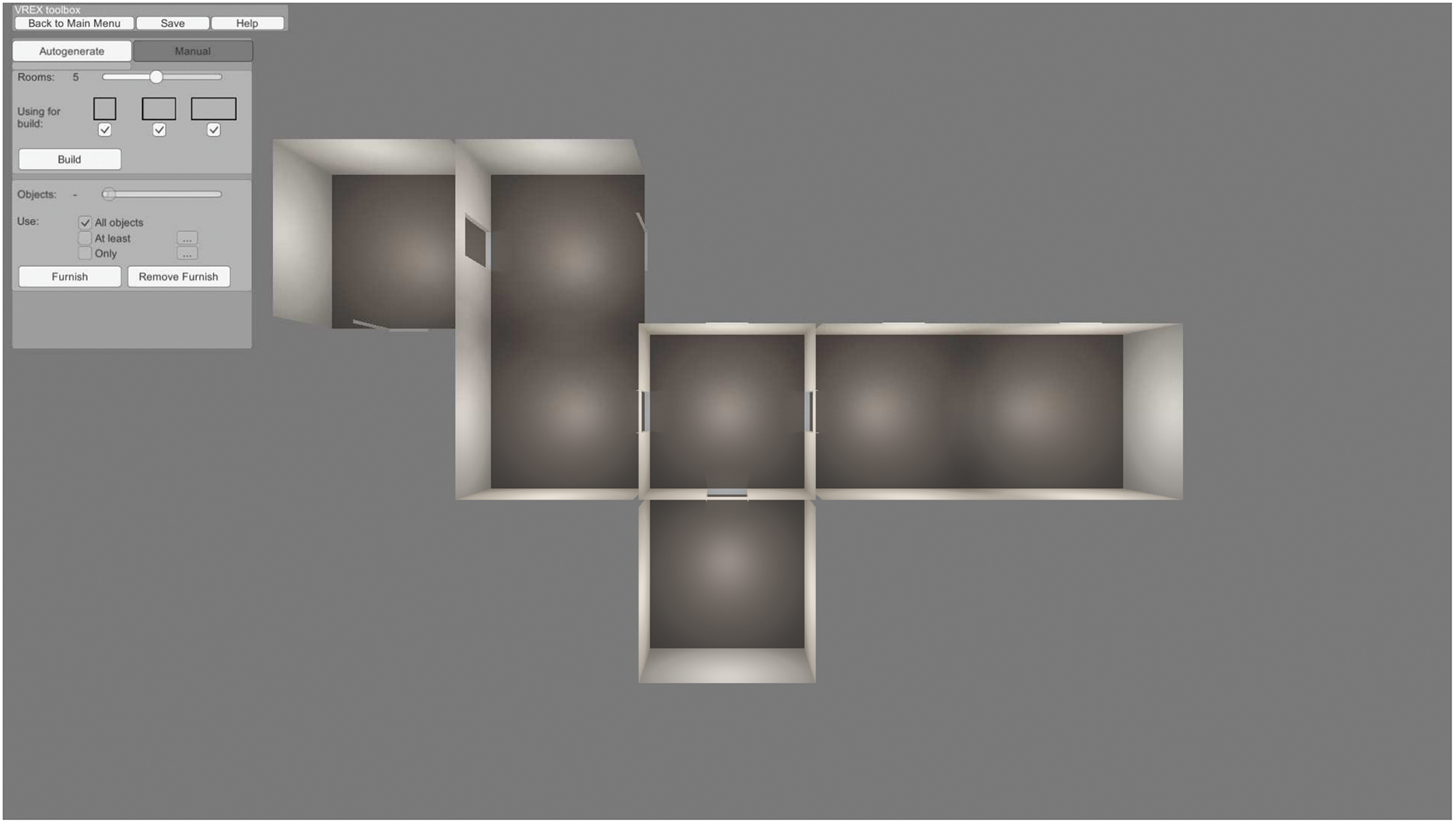
Sample environment generated procedurally with the parameters set to include five rooms with three possible sizes.

## Furnishing environments

After an environment is created either procedurally or by hand, it can then be populated with available 3D objects either automatically or manually. For automatic furnishing there can be a set number of objects in each room. The objects have pre-defined properties that place them according to a general logic - tables and chairs are placed on the ground, small object lay on the tables and shelves are attached to the wall etc. Automatic furnishing of objects saves time and produces a novel room layout on every instance. Figure 4 shows different autogenerated layouts for the same environment.

**Figure 4.**
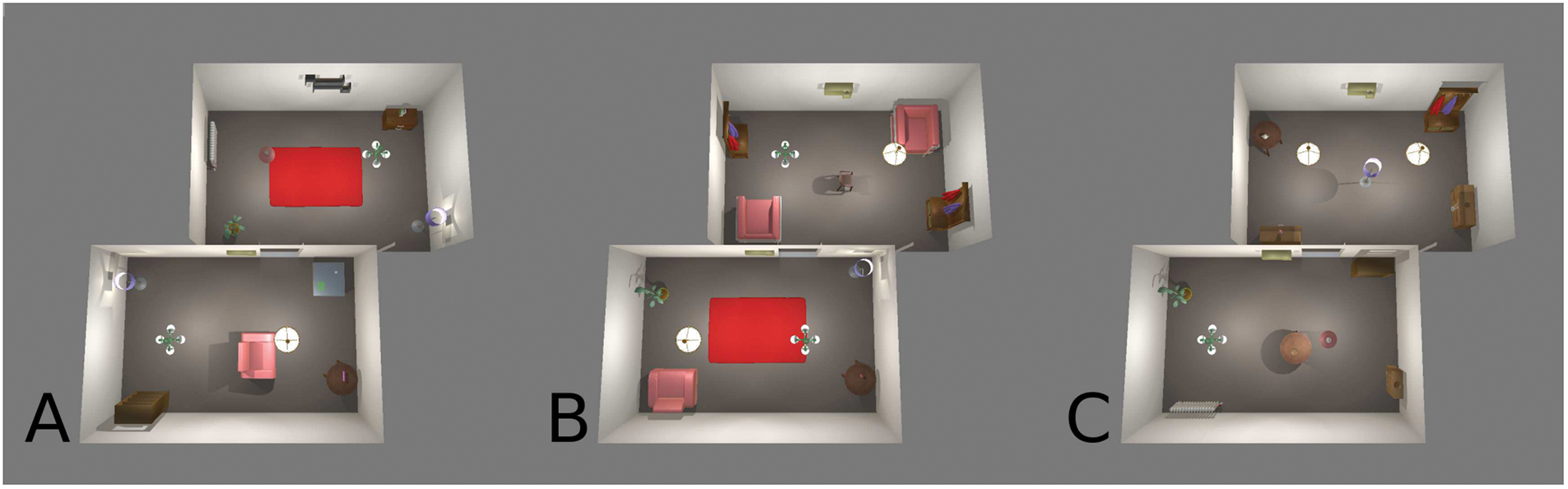
Three different results with the automatic furnishing option (A, B & C)

There are a number of openly licenced objects available in the toolbox (figure 5). With random placement some objects may end up in illogical positions from an interior design point of view. This can be easily corrected in the 3D editor.

**Figure 5.**
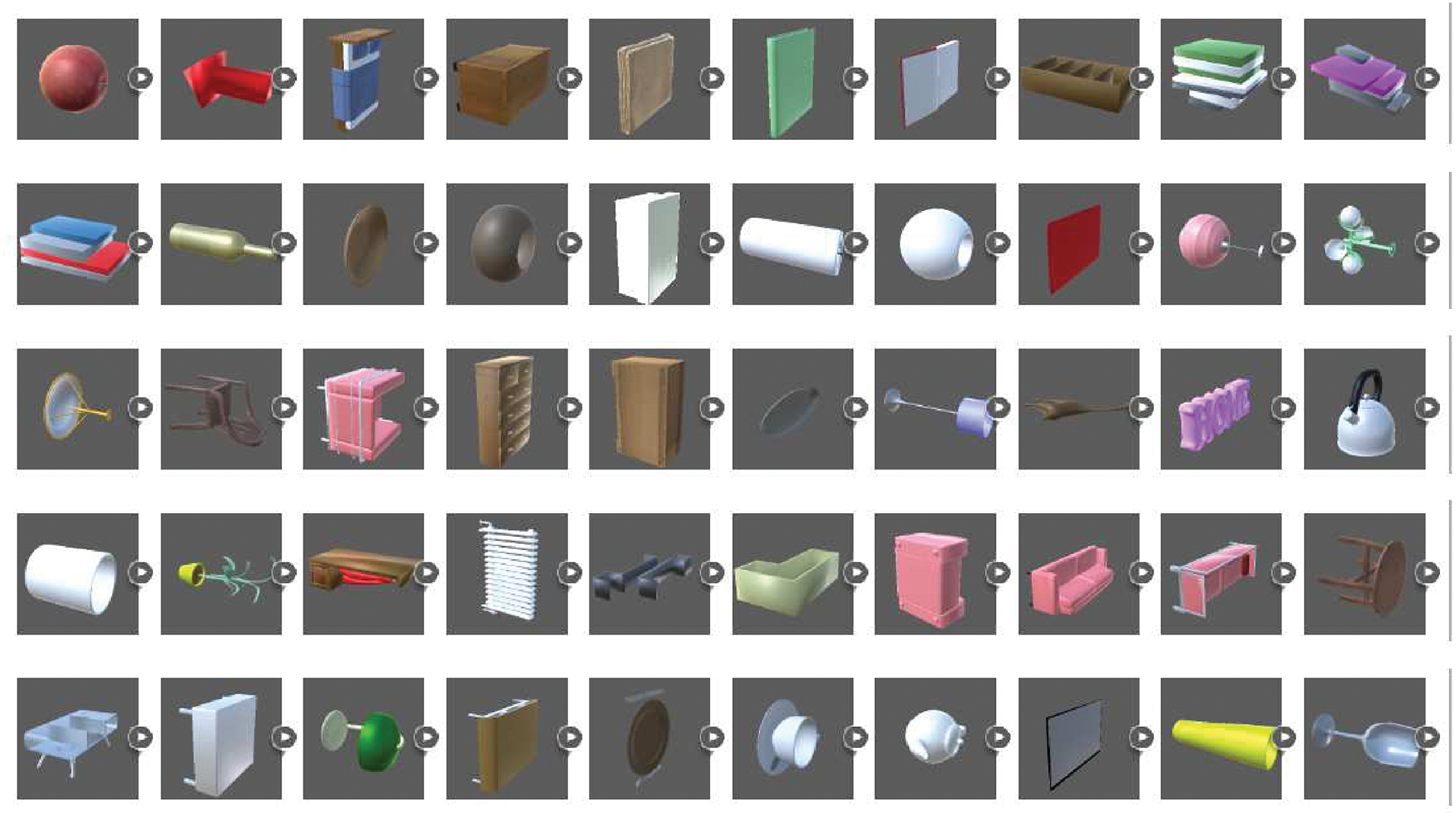
An overview of the available pre-made objects in VREX.

## 3D editor

For fine-tuned control VREX comes with an editor where all the objects in the room can be manually adjusted in the 3D world or new objects added (figure 6). Navigating the 3D editor is achieved with the mouse and keyboard. All objects in the scene can be moved, rotated and scaled, and the diffuse colour can be changed. Here the user can also define experiment-specific behaviours when an object is selected.

**Figure 6.**
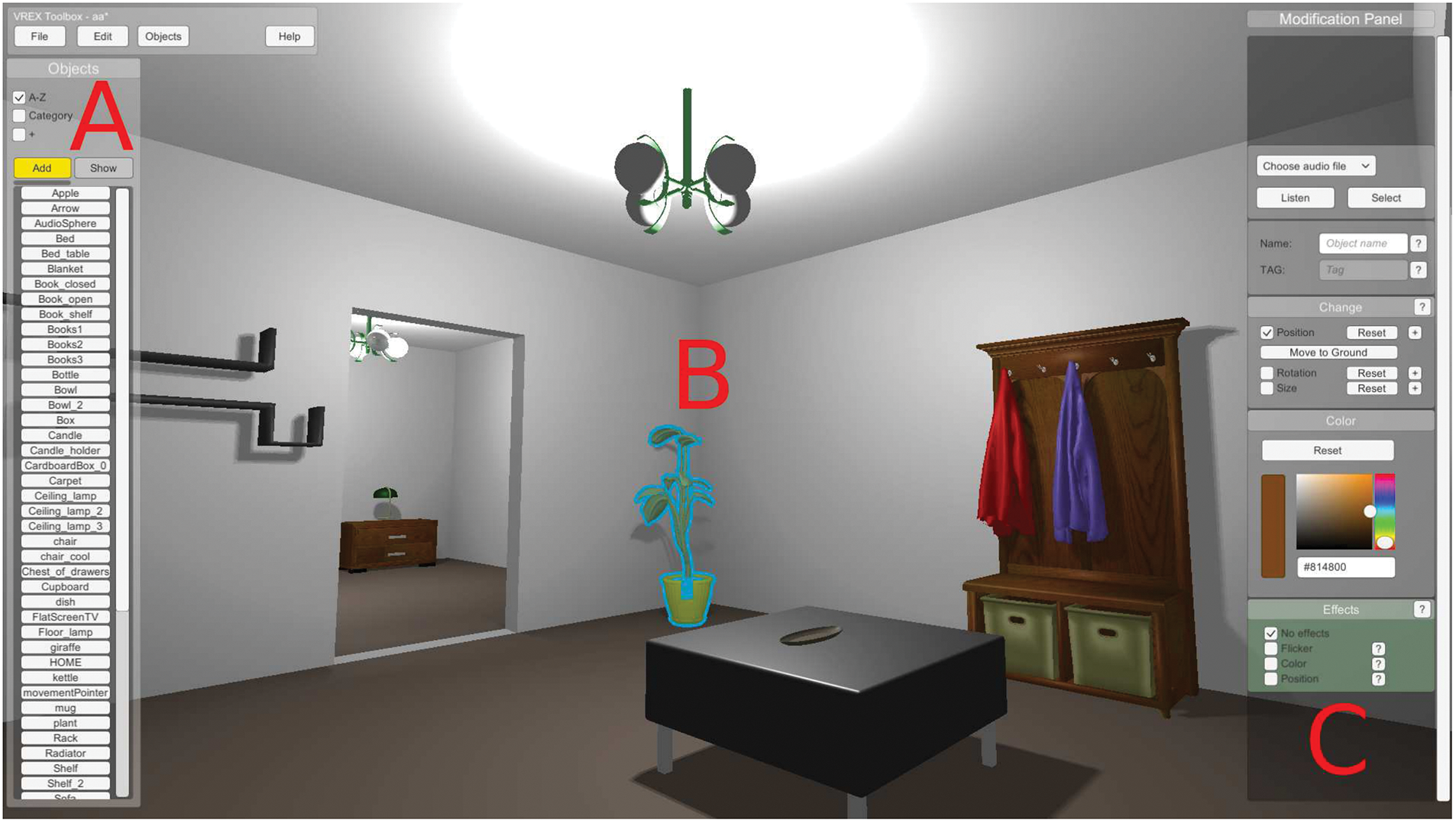
The 3D editor allows to navigate and place individual objects in the scene manually (A), select objects in the environment (B) and change different properties of the chosen object, such as position, rotation, size and colour (C).

## Experiment-specific behaviours

VREX currently supports two experimental paradigms - change blindness and false memory. Objects marked in the change blindness experiment can be modified whenever the object falls outside the field of view of the VR headset. The researcher can choose to change the object’s visibility, colour, or location (figure 6C). The chosen appearance change will alternate between two states. After identifying the change in an experiment, the participant can click the response button, which logs the response time. A cursor then appears in the centre of view in the response phase to aid selection. The participant can point at the suspected object with the centre of view of the headset and clicking the response button again will save the answer. After that the next level is automatically loaded.

In false memory related experiments VREX supports logging all the objects seen by the participant during a trial and later modifying their position in a room or presenting them for cued recall. Recall only contains items seen by the participant and optionally distractor items chosen by the experimenter. Recall can also have a set time limit. VREX currently supports two modes of recall - objects are either placed in an empty field or shown one by one. The participant must select all previously seen objects by moving close to the object and pressing the response key or answering yes/no in the case of one by one presentations.

## Creating experiments

After the environments have been finalized, experiment creation can begin. A new experiment must have a type, either change blindness or false memory. Each type has a specific set of options available regarding the time limits and test levels. Next the researcher can sequence all necessary environments in an ordered or randomized groups, add instructions to the participant and set appropriate test conditions.

## Additional features

## Adding custom models

Some experiments call for specific 2D or 3D objects that are not available in the standard VREX project library. Bringing in custom models involves using the standard Unity import pipeline and creating prefabs with specific properties outlined in the user manual. Prefab objects must have a tagged interconnector component attached and an origin point at the base of the geometry. For optimal performance, objects should not have an excessive polygon count. There is also a tutorial video detailing the whole process here: https://youtu.be/6YvTJsYvkxc

## Spatial audio integration

Spatial audio improves immersion in virtual reality [6]. The toolbox allows for easy placement of 3D sounds within the environments through the 3D editor, without using the standard Unity menus (figure 6, upper right). The implementation uses Unity built-in spatial audio algorithms and head-related transfer functions to create realistic sound transformations. For performance reasons it is advised to keep the total number of 3D sounds low.

## Four different locomotion systems

In most experiments the participant must be able to move around large virtual environments in conditions where the real space is limited. There are options in the Unity inspector window to choose the locomotion system for a given experiment. This can be either regular movement with the gamepad or keyboard, teleportation between points (using the default “j” key) or incremental movement and turning. There is also an option to automatically move the participant on a previously defined path. This option can be set up in the 3D editor.

## Data logging and timing

Every time an experiment is run through VREX, a data log is created in the VREX system folder under Results. The following is logged by default: Participant ID, start time of the experiment, current environment, environment order, test type, score (correct or false answer) and elapsed time. In case of change blindness and memory experiments additional parameters are stored, such as change count, changed object(s) and participants answer(s). Time is measured with millisecond precision.

## Template experiment for custom developments

A more experienced user may want to modify or extend the functionalities of VREX. A collection of template scripts with basic elements in place to develop a custom experiment with new behaviours can be found under the assets folder in the Unity project. For example it is possible to create a script for tracking the user’s trajectory for spatial navigation tasks inside the environments, change the input scheme, add parameters that should be logged during an experiment or tweak the user interface for the participant.

## Compatibility and extensibility

Different labs have different setups. The toolbox currently only works with the Oculus Rift DK2 and CV1 headsets due to the way the virtual camera is implemented. Other peripherals such as Leap Motion hand tracker or the HTC Vive VR headset can be introduced via the standard Unity interface.

## Use Cases

VREX is designed to allow easy creation of indoor VR environments and specific experiments. Although the software is far from being complete, it can already be used for practical research by novice programmers. Detailed examples of both change blindness and false memory experiment types are described below.

## Example 1 - building a change blindness experiment

Change blindness occurs when a person is unable to notice a big change in a scene, unless the scenario is presented numerous times or a hint is given [15]. With VR we wanted to study the effects of 3D depth on change blindness performance. Conducting such a study with the VREX toolbox is easy and requires no coding. First, we create ten virtual environments, procedurally and manually choosing and placing objects at different 3D depths from the participants initial position. We then hand-pick the objects that will be set to exhibit the desired change blindness behaviour. We chose the appearance/disappearance, meaning that certain objects change visibility whenever they are outside the field of view of the headset. All the environments are ordered randomly in the experiment menu, preceded with a text to display instructions to the participant. In each environment the participant has a set amount of time to look around, trying to spot the change that happen whenever he/she is looking away from the changing object. When the time is up, the participant chooses the object she thinks was changing by looking at its general direction and pressing the response key on a gamepad. An overview of the experiment from the participants’ eyes can be seen on figure 7. A formal experiment following these principles has demonstrated that such changes take a long time to be spotted and that changes further away in depth (i.e. in the 3D background) take longer time to be detected [16].

**Figure 7.**
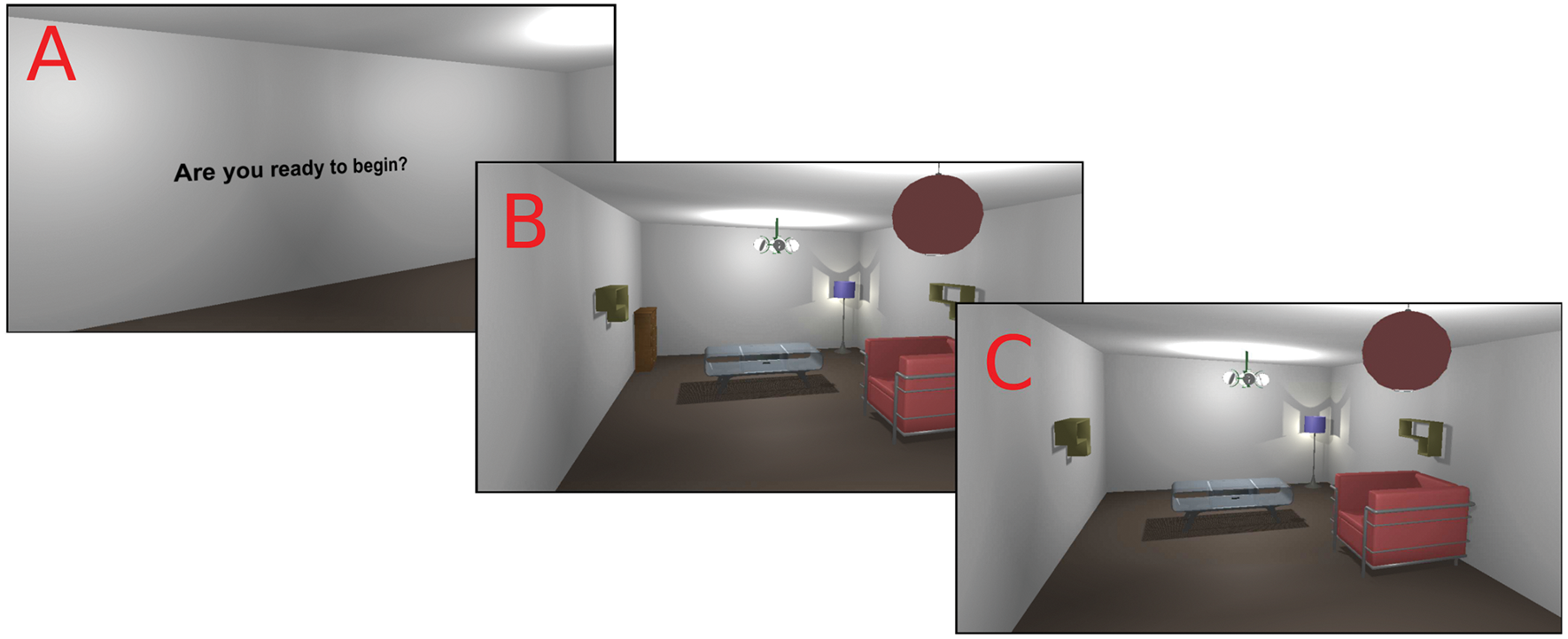
An overview of the change blindness experiment from the participants view. First the player is welcomed with a text message (A) and then transported to the experimental environment (B). While looking around in the room, the cupboard changes its visibility when out of the headsets field of view (C).

## Example 2 - building a false memory experiment

False memory occurs when an individual remembers events that did not happen or were different in reality [17]. Studying false memory with VREX is again greatly simplified with a built in memory experiment template and a clear menu system. For example we can study if participants mistakenly remember seeing objects in the environment that were not present. First we create the environment by generating a series of connected rooms placed procedurally and then automatically populate the rooms with objects. In the experiment window we determine the study design, assign text blocks (e.g. instructions to the user) and test procedures (e.g. yes/no recognition test). Once the participant has experienced every virtual room in the experiment, they are “teleported” to the recognition test level and asked if they recall seeing different objects one by one. Some objects previously seen by the participant are automatically replaced with distractors by the toolbox. The memory performance of the participant is assessed according to signal detection theory by counting true hits (answering “yes” on objects that were presented), false alarms (answering “yes” to objects not presented), correct rejections (answering “no” to objects not presented) and misses (answering “no” to objects that were present). The participants answers are recorded and logged, so further data analysis can take place outside of VREX.

## Limitations

VREX user interface is designed with very specific tasks in mind. This means that many options are limited and the user can not access the full potential of Unity through the toolbox menus. For general ease of use all key bindings are currently hardcoded and can not be easily remapped by the user, and this could be problematic for some types of experimental plans. Convenient access to such settings is where MazeSuite and WorldViz Vizard currently outperform VREX.

Another limitation is that as VREX relies heavily on Unity 5, major future updates to the game engine may cause compatibility errors. This is common for all add-ons and recommended Unity version numbers should be noted when using the toolbox. For the standalone version of VREX this is not an issue.

## Conclusions

We have presented VREX, a Unity toolbox for VR research in psychology. By adding many powerful features and a simple user interface, the tool is designed to empower researchers with no or little prior knowledge in game engines to start working with VR. Together with the community we aim to develop the application further. The website of the VREX toolbox can be found on the following link: vrex.mozello.com

## Declarations

### Ethics approval and consent to participate

Not applicable

### Consent for publication

Not applicable

### Availability of data and materials

Project name: VREX: A UNITY^®^ TOOLBOX FOR VR EXPERIMENTS

Project home page: http://vrex.mozello.com/

Archived version: https://drive.google.com/open?id=0B97-aac1_IQ5eEdiaW83TkZsa2c

Operating system: Microsoft Windows

Programming language: C#

Other requirements: Unity Engine 5 (none for standalone version)

License: Creative Commons BY-SA licence

Any restrictions to use by non-academics: none

### Authors' contributions

MV, TK, RV & JA designed the general principles of the toolbox. MV created the visual elements. TK designed the user interface. MK & KK programmed the main functionalities. VS & MM programmed additional functionalities. MV & JA wrote the manuscript. All authors read and approved the final manuscript.

## Acknowledgements

The authors would like to thank Kristiina Kompus, Andero Uusberg and Helen Uusberg for early input; Egon Elbre and Ardi Tampuu for consulting on how to build the result database; Iiris Tuvi, Liisi Kööts-Ausmees and Kristiina Kompus for alpha testing; Computational Neuroscience Lab and Department of Psychology of the University of Tartu.

